# Transcriptome wide analysis of natural antisense transcripts shows their potential role in breast cancer

**DOI:** 10.1101/176164

**Authors:** Stephane Wenric, Sonia ElGuendi, Jean-Hubert Caberg, Warda Bezzaou, Corinne Fasquelle, Benoit Charloteaux, Latifa Karim, Benoit Hennuy, Pièrre Freres, Joëlle Collignon, Meriem Boukerroucha, Hélène Schroeder, Fabrice Olivier, Véronique Jossa, Guy Jerusalem, Claire Josse, Vincent Bours

## Abstract

Non-coding RNAs (ncRNA) represent at least 1/5 of the mammalian transcript amount, and about 90% of the genome length is actively transcribed. Many ncRNAs have been demonstrated to play a role in cancer. Among them, natural antisense transcripts (NAT) are RNA sequences which are complementary and overlapping to those of protein-coding transcripts (PCT). NATs were punctually described as regulating gene expression, and are expected to act more frequently in *cis* than other ncRNAs that commonly function in *trans*. In this work, 22 breast cancers expressing estrogen receptors and their paired healthy tissues were analyzed by strand-specific RNA sequencing. To highlight the potential role of NATs in gene regulations occurring in breast cancer, three different gene extraction methods were used: differential expression analysis of NATs between tumor and healthy tissues, differential correlation analysis of paired NAT/PCT between tumor and healthy tissues, and NAT/PCT read count ratio variation between tumor and healthy tissues. Each of these methods yielded lists of NAT/PCT pairs that were demonstrated to be enriched in survival-associated genes on an independent cohort (TCGA). This work allows to highlight NAT lists that display a strong potential to affect the expression of genes involved in the breast cancer pathology.

## Introduction

Over the past decade, RNA sequencing technology allowed to discover that the non-coding part of the genome represents around 1/5 of all transcript amount ^1,2^. These non-coding RNAs (ncRNA) are less conserved between species than protein coding genes, but more than introns and random intergenic regions ^3,4^. It is therefore likely that these non-coding transcripts have biological roles, which are being progressively deciphered, but still remain largely unknown. Around 30-50% of the protein coding gene loci are additionally expressing ncRNA in an opposite direction of the protein coding gene ^4,5^. These naturally occurring antisense transcripts, called NATs, have been less studied than the other classes of ncRNA for technical reasons because their detection and quantification require to preserve information about the transcript originating strand along the sequencing process. Indeed, standard RNA sequencing and expression micro-array techniques require double-stranded cDNA synthesis, which erases RNA strand information, leading to an expression quantification that is the sum of the expressions of the coding RNA and its corresponding NAT. Commercial kits allowing to gather this information have only been made available recently, paving the way to high-throughput studies of stranded RNA sequencing.

NAT expression is subjected to the same expression regulation than other genes, but NATs accumulate preferentially into the nucleus - associated to chromatin - unlike coding mRNAs which are located into the cytoplasm. NATs are also found in other cellular compartments such as mitochondria ^6,7^. NAT expression is described in many punctual examples to affect the activity of their sense or neighboring genes in biological events like cell differentiation and carcinogenesis, distinct molecular mechanisms being involved ^8–11^. NATs can regulate gene expression in *trans* or in *cis.* Given the fact that both the sense and antisense transcripts are transcribed from the same genomic region, it is expected that antisense transcripts behave more frequently in *cis* than other ncRNAs that commonly function in *trans* ^11^.This last feature means that NATs may regulate their protein coding gene counterpart at the same locus, which is of great interest from the therapeutic point of view: NATs may thus provide a unique entry point for therapeutic intervention on targeted genes by the use of ASO (antisense oligonucleotides) that are drugs already FDA-approved for several diseases ^8,12–14^.

To date, a few studies have been performed at the whole transcriptome scale to investigate the role of NATs in the context of breast cancers. These studies have demonstrated that pairs of NAT/PCT are globally deregulated in this pathology ^15–17^. However, none of those studies compared the whole transcriptome of paired tumorous and healthy tissues of the same patients, with a technology that keeps the strand information of the transcripts. Yet, such an experimental design would be needed to explore if NAT tumor deregulations are cancer-specific, in order to define if they could play a role in the pathology. Here, we describe the results of such an experimental design, in a cohort of 22 ER+ breast cancer patients whose paired healthy and tumorous tissues were analyzed by stranded RNA sequencing.

This work allows clarifying the role played by NATs to regulate their protein coding gene counterpart on the same locus in the breast cancer pathology and to quantify to what extent this phenomenon is occurring. We first defined 3 lists of NAT/PCT pairs that are both deregulated between healthy/tumorous tissues and related to NAT-specific regulations. Next, we demonstrated that those lists are enriched with survival-associated genes. Finally, we established a list of breast cancer-related genes probably regulated by their NATs that could be targeted by ASO in a therapeutic objective.

## Results

The role played by NATs on the expression regulation of their corresponding coding transcripts is still largely unknown, as well as if this potential regulation could play a role in the ER+ breast cancer pathology. Our study experimental design used to answer this question at the whole genome scale is depicted in Figure 1. Twenty-two tumorous tissues of ER+ breast cancer patients, as well as their paired adjacent healthy tissues, were subjected to strand-specific RNA Sequencing and DNA copy number analysis by CGH array. The patient characteristics are summarized in Table 1. The cohort contains only tumors larger than 20mm, and is equally divided between luminal A and B sub-types, and between highly (Ki67>19%) and moderately (Ki67<19%) proliferating tumors. Most of them present a Bloom grade of 2 and 3.

**Figure 1.**
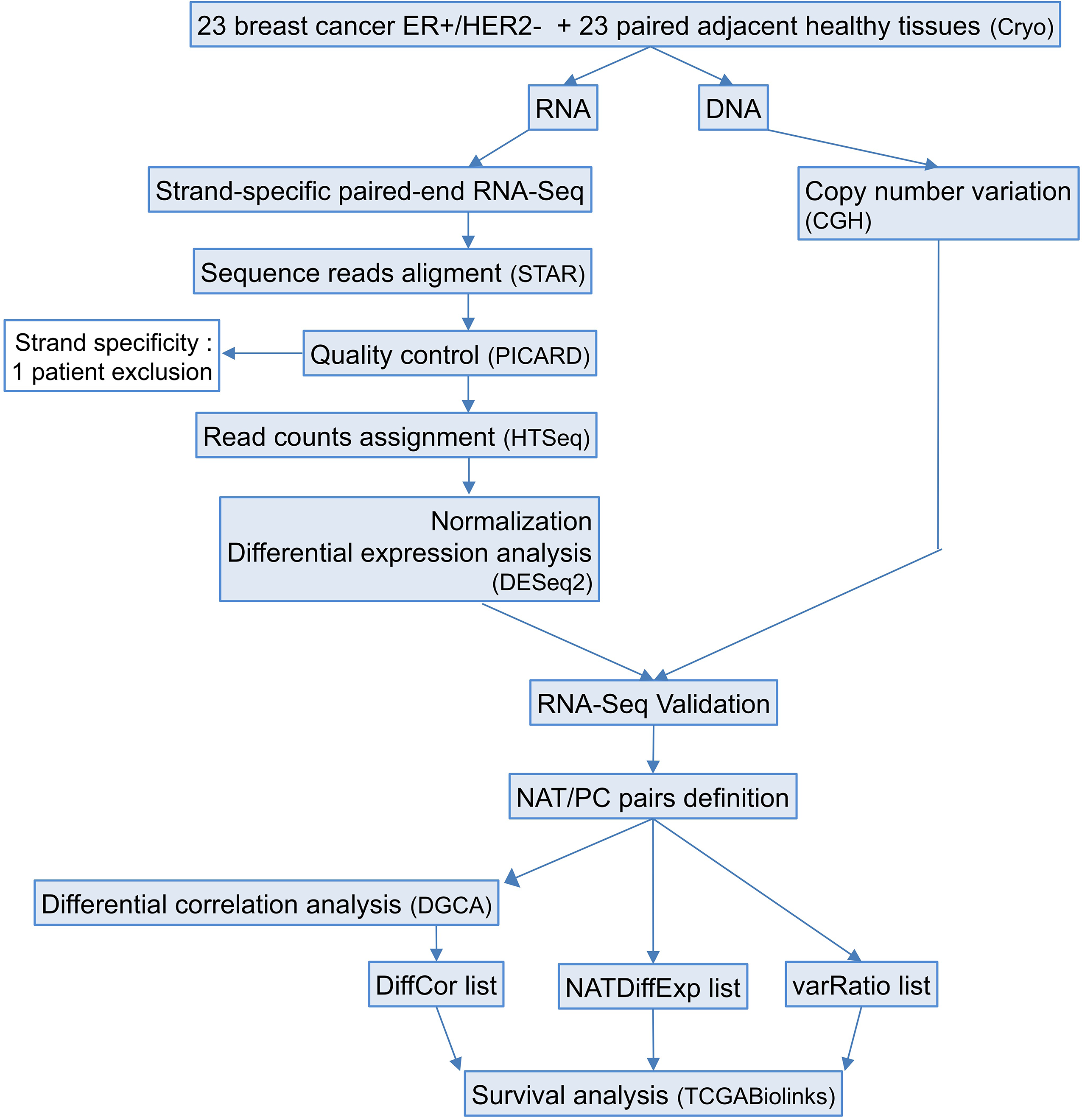
Study workflow. RNA and DNA were simultaneously extracted from 23 breast cancer ER+/HER2- tumors and their paired adjacent healthy tissues. Strand-specific paired-end RNA sequencing and comparative genomic hybridization (CGH) were performed. Quality control steps and RNA-Seq validation were performed and lead to the elimination of one patient because of a poor strand specificity of this sample. This strategy allowed to study differential expression of NATs and PCTs between tumors and healthy tissues, and to perform differential correlation analysis of NAT/PCT pairs. Three lists of genes with deregulated NAT expression in the tumors that could potentially affect their corresponding PC expression were extracted, and the coding genes they contain were subjected to survival analysis with an external cohort (TCGA).

**Table 1.**
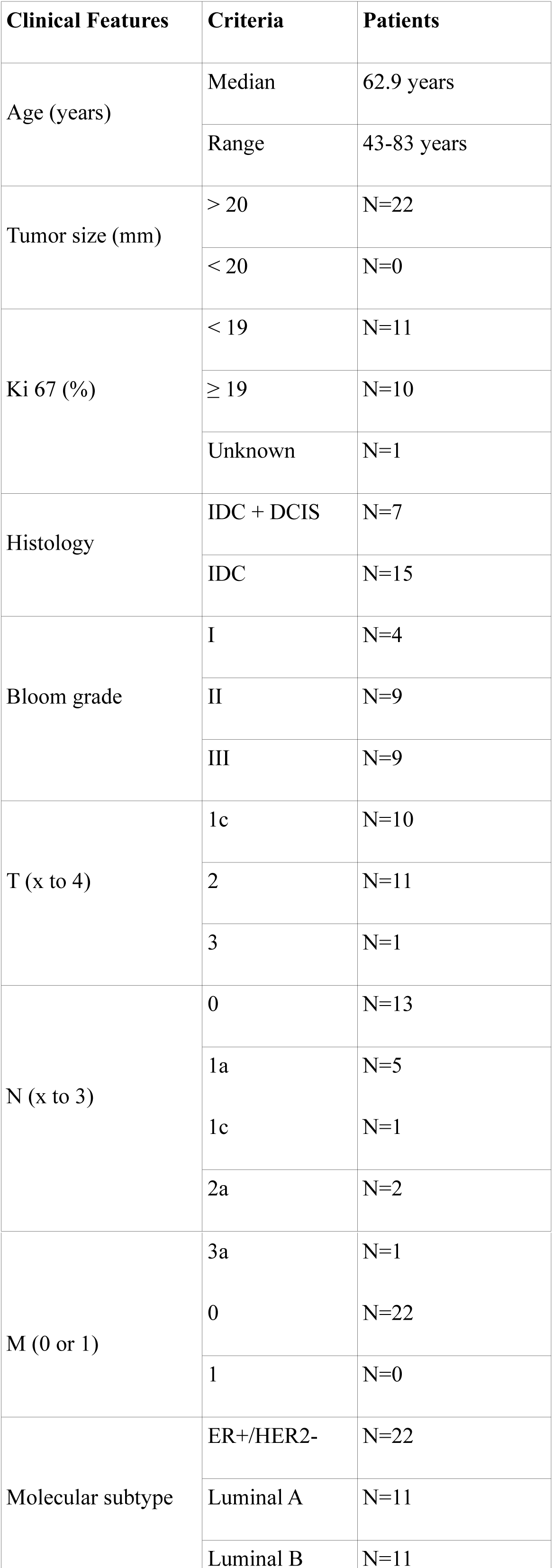
Patient clinicopathological characteristics

### RNA-Seq Validation

The combined analysis of the DNA copy number variations and modifications of RNA transcripts expression levels between tumor and healthy tissues validates our RNA-Seq analysis: the overall expression levels of coding gene transcripts inside genomic amplification or deletion newly acquired in the tumor were respectively increased and decreased, as expected (Figure 2 A).

**Figure 2.**
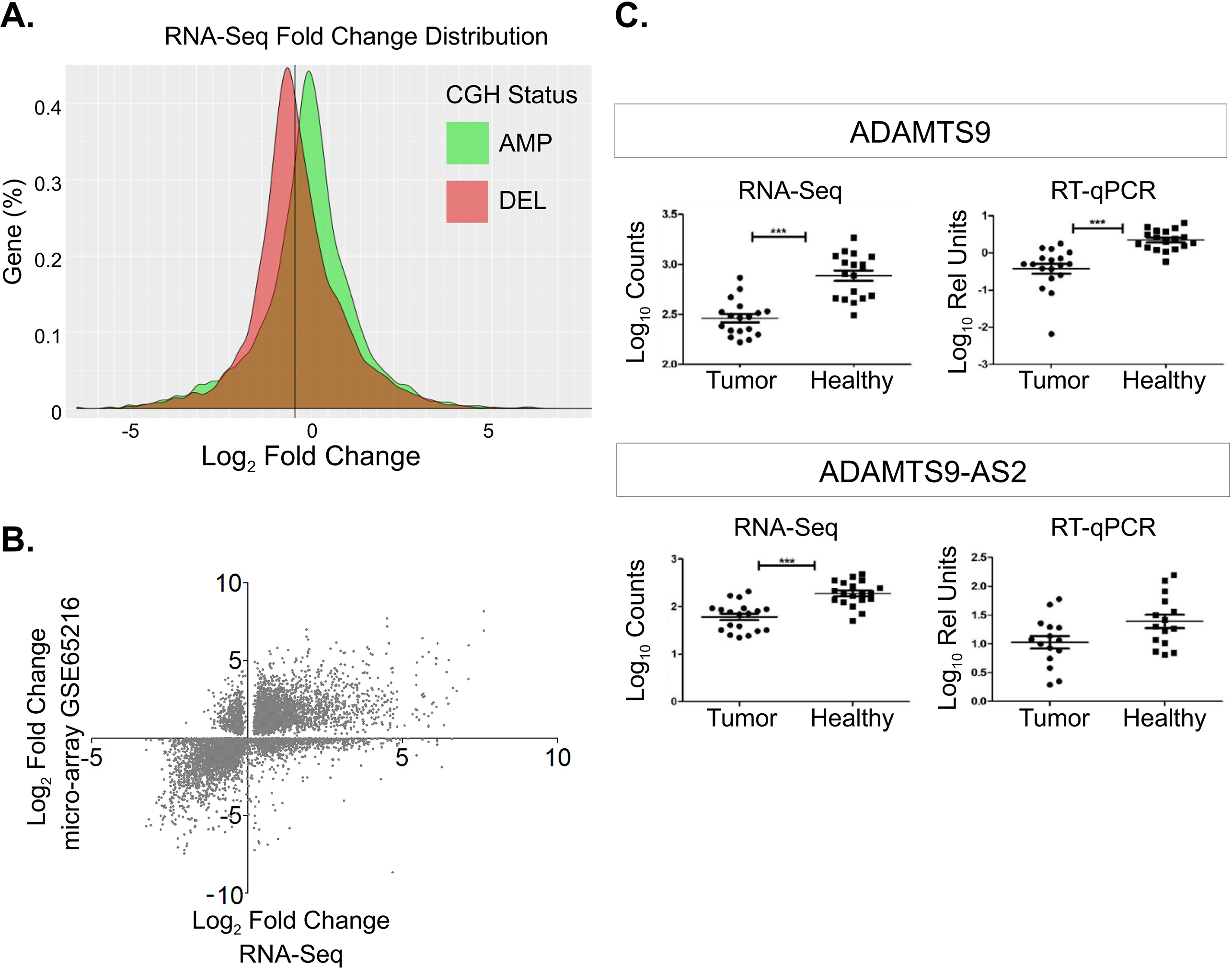
Validation of RNA-Seq. **A.** Fold change distributions of genes, as determined by RNA-Seq, which are located in somatic copy-number alterations (amplifications or deletions), as determined by CGH. The distinct curves show a clear effect of the copy-number alterations on the gene expression (fold-changes). As expected, genes located in genomic amplified regions in the tumor showed increased expression, and conversely. **B.** Gene expression fold-changes between tumor and healthy tissues obtained in the current RNA-Seq study were compared to those described in an external Affymetrix micro-array dataset GSE65216. This comparison showed a global concordance of the results, with a Spearman correlation coefficient of 0.613 (p-value < 0.001). **C.** The relative expression of the protein coding ADAMTS9 and its NAT, ADAMTS9-AS2, in tumors and healthy tissues obtained by RNA sequencing and by RT-PCR were compared. The RT-qPCR values were normalized by the expression of the endogenous control gene B2M. [p-value <0.001 (***)].

Moreover, the gene expression changes between healthy and tumor tissues obtained in our RNA-Seq dataset were compared with those obtained in an independent dataset (GSE65216). Gene expression variations between 10 mammary healthy tissues and 22 ER+ tumors (11 luminal A and 11 luminal B) were extracted using Geo2R ^18^. The expression fold-change of genes that were found to be differentially expressed with an adjusted p-value <0.05 between healthy and tumoral tissues in both our and the GSE65216 datasets were compared, and present an average Spearman correlation coefficient of 0.613 (p-value<0.001). Moreover, 76.6% of those genes were differentially modulated in the same direction (Figure 2 B). At a smaller scale, some RT-qPCR experiments were performed on the RNA samples that were used for our RNA-Seq study to confirm variations of several transcripts expression between tumor and healthy tissues. Among others, the downregulation in tumors of the ADAMTS9 tumor suppressor and its NAT, ADAMTS9-AS2, were confirmed by RT-qPCR (Figure 2 C).

### NAT expression accounts for 17% of their coding counterpart in healthy tissues and increases to 26% in tumors

We next defined the pairs of protein-coding genes and their corresponding antisenses as detailed in the material and methods section. This list can be found in Supplementary Table and contains 9632 NAT/PCT pairs where at least one patient has a non-null expression for PC or AS, either in normal tissue or in tumor. As 19846 coding transcripts were expressed in mammary tissues, 49% of coding transcripts have a concomitant corresponding NAT expression. Globally, NAT read counts represent 17% of their coding counterparts in healthy tissues and 26% in tumors (Table 2), suggesting a global increase of the expression levels of NATs in mammary tumors. Moreover, the average read counts ratio between PCT/NAT gene pairs expressed simultaneously by a locus is 1544 in healthy tissues and 1013 in tumors (Supplementary Table).

**Table 2:**
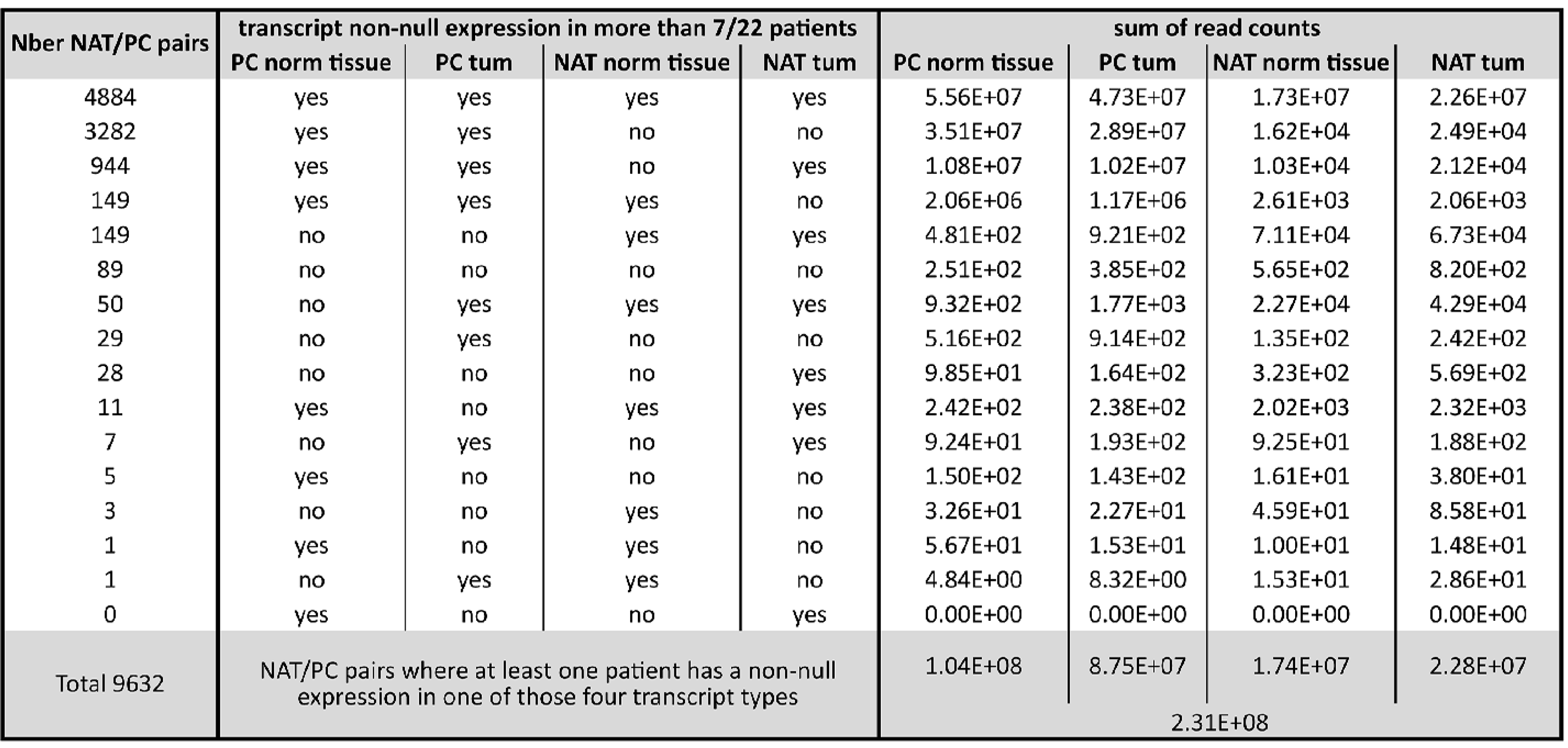
Distribution of the relative expression intensities of NAT and their corresponding PC among the 9632 NAT/PC pairs. This study was focused on NAT/PCT pairs where both the PCT and the AS were expressed in at least 7 out of the 22 patients, both in the tumor and the healthy tissue. This group of 4884 gene pairs contains 60% of the total reads counts, and the NAT/PCT ratio expression is increased in tumors.

Unexpectedly, we observe that the fold change distributions of NATs present both in genomic amplifications and deletions are shifted towards higher fold-changes than the corresponding distributions based on protein-coding genes (Figure 3 A).

**Figure 3.**
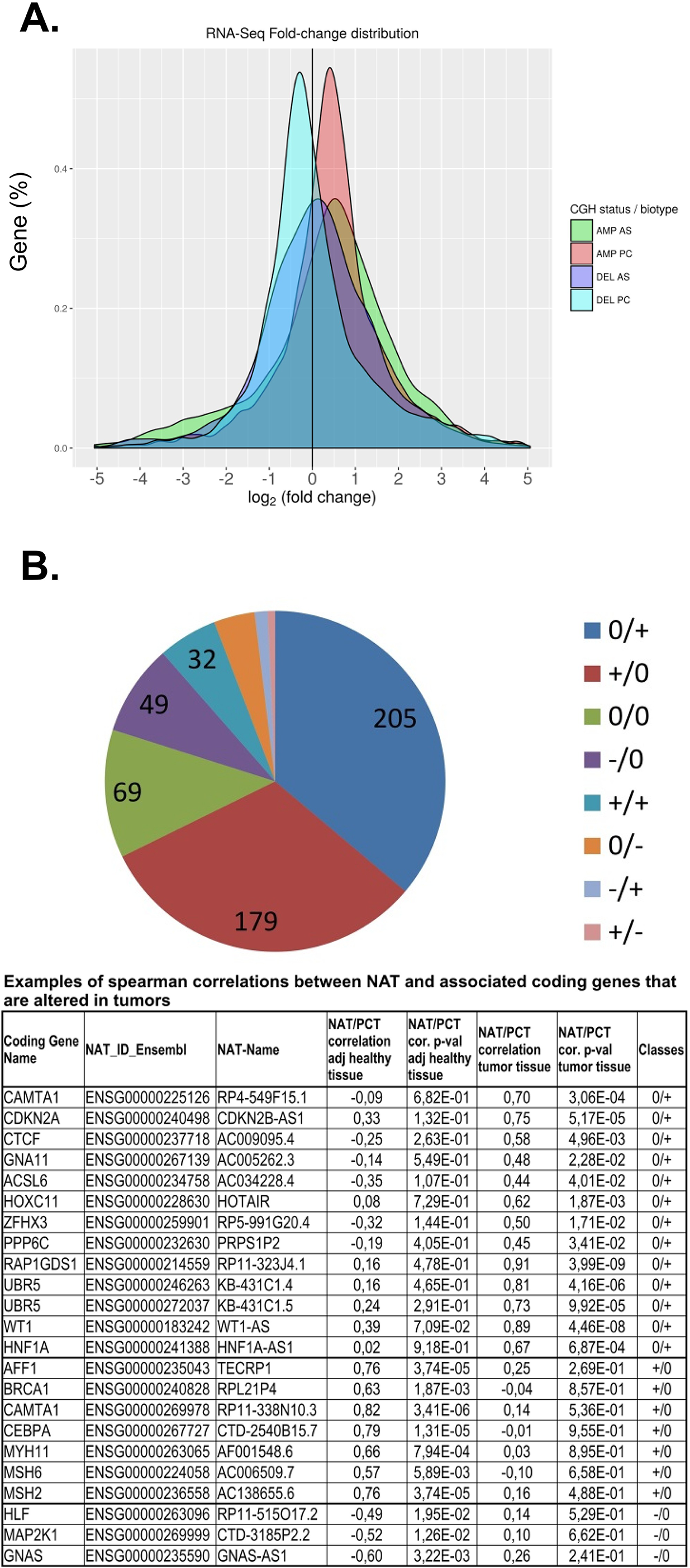
A. Fold-change distributions of NAT present in genomic amplifications and deletions. NAT expression values are shifted towards higher fold-changes than the corresponding distributions of their protein coding genes. **B. Schematic representation of the proportion of the different classes of differential correlations between tumors and healthy tissues.** Mainly positive correlations of expression between NAT and their corresponding PCT are created, or lost in tumorous tissues. The numbers indicated in the graph are the numbers of NAT/PCT pairs in this category; +/+ = significant positive correlation between NAT and PCT exists in the adjacent healthy tissue and is conserved in the tumor; +/- = significant positive correlation between NAT and PCT exists in the adjacent healthy tissue and becomes negative in the tumor; +/0 = significant positive correlation between NAT and PCT exists in the adjacent healthy tissue and is lost in the tumor; -/- = significant negative correlation between NAT and PCT exists in the adjacent healthy tissue and is conserved in the tumor; -/+ = significant negative correlation between NAT and PCT exists in the adjacent healthy tissue and becomes positive in the tumor; -/0 = significant negative correlation between NAT and PCT exists in the adjacent healthy tissue and is lost in the tumor; 0/0 = no significant correlation between NAT and PCT exist, nor in the adjacent healthy tissue nor in the tumor.

Based on the levels of expression in the different tissue types, we chose to focus on NAT/PCT pairs where both the PC and the AS were expressed in at least 7 out of the 22 patients, both in the tumor and the healthy tissue. This represents more than 60% of the total read counts of NAT/PCT pairs (Table 2). In this group of 4884 genes pairs, NAT expression is greatly increased and accounts for 31% of their coding counterpart in healthy tissues and 47.8% in tumor. This gene sub-group contains PC genes that display the stronger potential of being regulated by their NAT counterparts.

### Positive correlation of expression between NAT and their corresponding PCT are created in tumorous tissues

In order to highlight newly appearing or disappearing correlations between NATs and their corresponding PCTs in the tumor, differential correlation analysis between all pairs of PCTs and NATs was performed with the DGCA software (v. 1.0.1) ^19^. Complete results can be found in Supplementary Table, showing a global positive correlation of expression between NATs and their corresponding PCTs: the mean Spearman correlation coefficients are 0.431 and 0.533 respectively in healthy tissues and tumors when a significant correlation is observed (p-value < 0.05), namely in 20% of the NAT/PCT pairs. The number of significantly correlated NAT/PCT pairs does not differ in healthy and tumorous tissues. A positive mean z-score (0.460) is observed in case of significant differential correlation of NAT/PCT between tumor and healthy tissues (p-value < 0.05), meaning that globally, in the 11% of NAT/PCT pair correlations that are deregulated in tumors when compared to healthy tissues (567/4884 pairs), the correlations become more positive. The proportion of the different classes of differential correlations between tumors and healthy tissues is depicted in Figure 3 B, highlighting the fact that mainly positive correlations of expression between NATs and their corresponding PCTs are created, or lost in tumorous tissues. Very few inversions of correlation were observed. Some examples of NAT/PCT pairs presenting deregulated correlation of expression in the tumor tissue are presented in Table 3 showing that genes that are well known in the breast cancer field are displaying deregulated correlation of expression with their antisense transcripts.

**Table 3:**
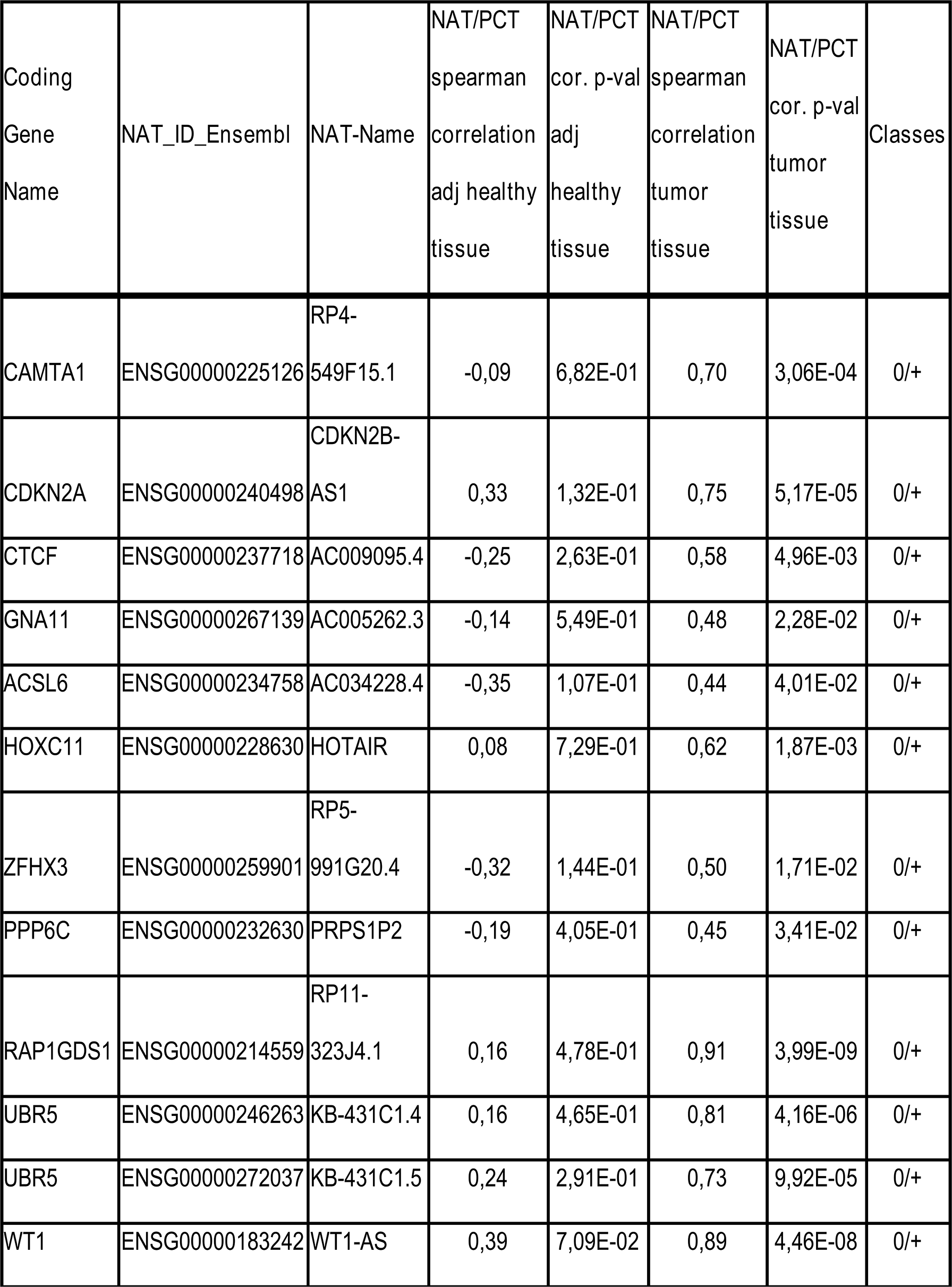
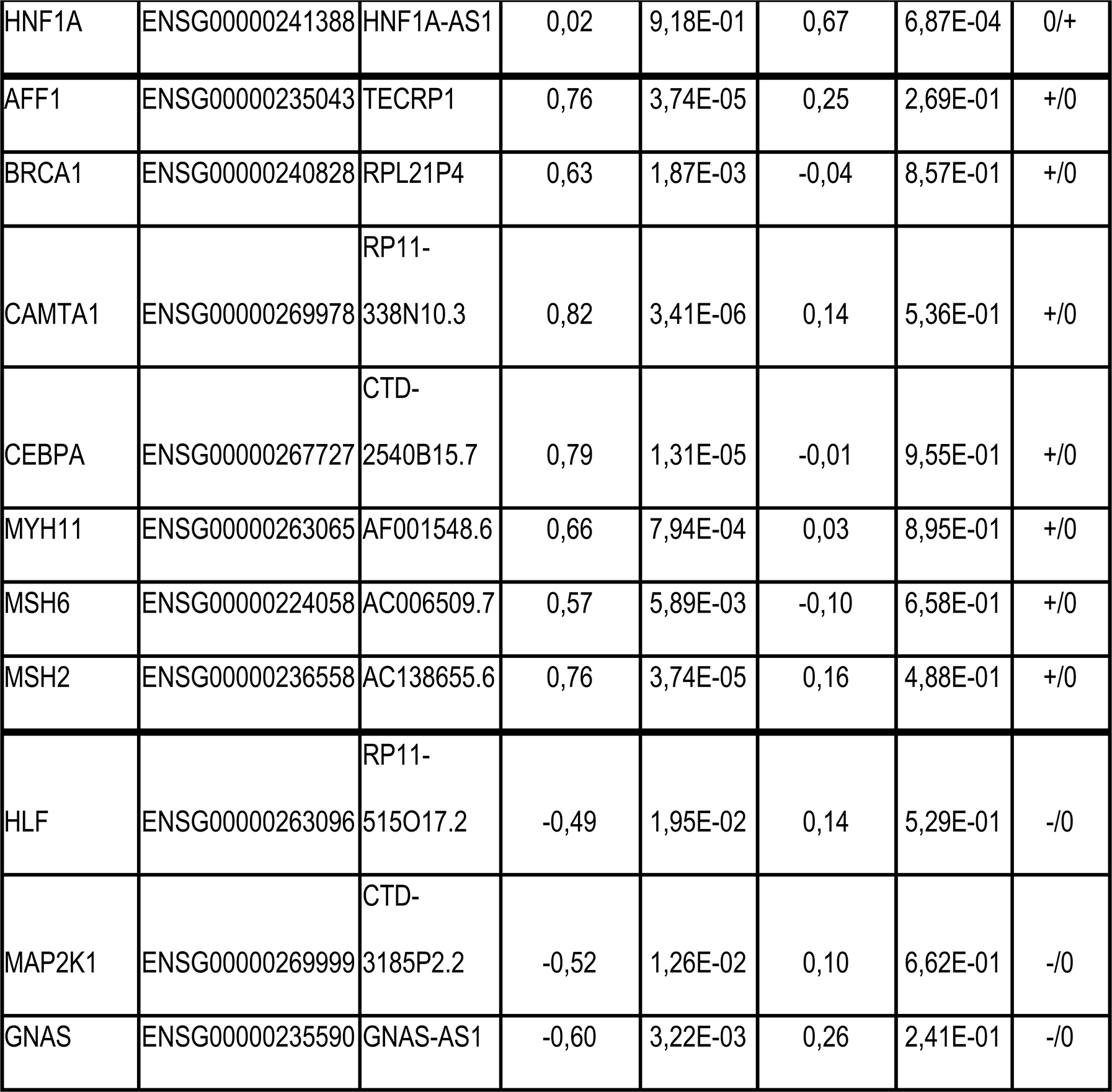
Examples of spearman correlations between NAT and cancer-associated coding genes that are altered in tumors when compared to adjacent healthy tissues.

### Protein coding genes exhibiting deregulated corresponding NAT expression in tumors are preferentially related to survival of breast cancer patients

Three different gene selection methods were used to extract NAT/PCT pairs potentially related to the breast cancer pathology out of the 4884 pairs.

Firstly, the previously described DiffCor is based on the differential correlation of NAT/PCT read counts between healthy and tumors tissues. A list of NAT/PCT pairs whose correlation is significantly different between normal and tumor tissues (p-value < 0.05) and whose correlation class differs between tumor and normal tissues (ie. 0/0, +/+, -/- classes are removed) has been selected and contains 441 NAT/PCT pairs.

The second method is based on the differential expression of the NATs between tumors and healthy tissues. A list of 738 NAT/PCT pairs where the NATs were significantly differentially expressed (adjusted p-value < 0.05) between normal and tumor tissues has been determined.

The third method is based on the variation of the NAT/PCT ratio between healthy and tumor tissues, and allows to define a third list, called VarRatio, which contains NAT/PCT pairs with extreme values on the distribution of the VarRatio (S8 – Supplementary Figures). This VarRatio list can be subdivided in leftmost and rightmost parts. Leftmost, 610 NAT/PCT pairs have a PCT/NAT ratio that decreases in the tumor either because of a down-regulation of the PCT expression, or an up-regulation of the NAT expression or both; and the reverse is observed for the 540 NAT/PCT pairs on the rightmost part of the distribution.

The 3 lists of genes can be found in the Supplementary Figures (S9 to S12) and as expected, many NAT/PCT pairs appear in several of those lists, which contains a total of 1784 unique NAT/PCT pairs deregulated in breast cancers.

To ascertain if the protein coding genes of the DiffCor, NATDiffExp, and VarRatio lists are implicated in the breast cancer pathology, their association with survival was computed based on the RNA-Seq samples from the TCGA dataset. Each of these three lists present a proportion of genes associated with survival in the TCGA dataset greater than the proportion obtained in a list of randomly chosen protein coding genes (Table 4), meaning that PCT exhibiting deregulated corresponding NAT expression in tumor are enriched in genes related to survival of breast cancer patients. A Pearson’s chi-squared test yielded statistically significant p-values for each of the 3 lists when compared to a list of randomly chosen protein coding genes.

**Table 4:**
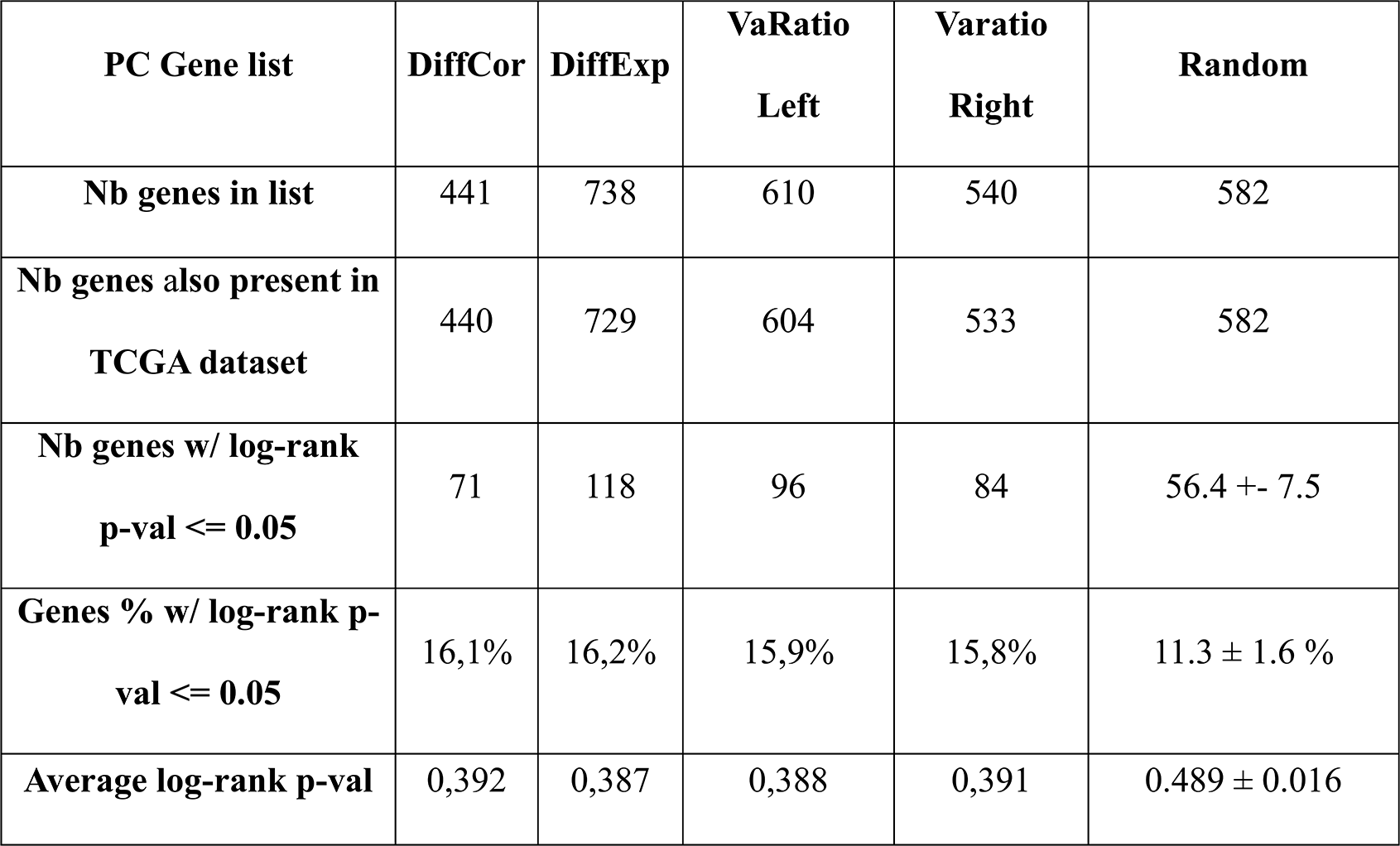
PC genes exhibiting deregulated corresponding NAT expression in tumor are preferentially related to survival of breast cancer patients. Protein coding genes of the 3 gene lists DiffCor, DiffExp and VarRatio (left and right) were tested for association with survival by means of a TCGA RNA-Seq dataset of breast cancers. The percentage of genes associated with survival in each list has been compared with randomly selected protein coding genes.

### 72 cancer genes present a deregulated profile of NAT expression in breast cancer samples

When the Cancer Gene Census list of genes from the COSMIC database (http://cancer.sanger.ac.uk/census) was compared with the content of our 3 lists of genes that are probably regulated by their NATs and implicated in the breast cancer pathology, 72 genes were found in common (S13 – Supplementary Figures). This list contains cancer genes that could be targeted by ASO, designed to interact with the corresponding NATs of those genes, in order to specifically regulate their expression.

## Discussion

Breast cancer constitutes a public health problem: around 1out of 8 women will suffer from it during their lifetime in industrialized countries. The most frequent subtype is the estrogen receptor expressing breast cancer (ER+/HER2-), with 75% of occurrences. In case of primary disease, most patients are treated by surgery with or without radiotherapy and endocrine therapy. However, a large number of those cancers will suffer from a relapse and develop metastases - a major life-threatening event which is strongly associated with poor outcome - and require chemotherapy in case of symptomatic visceral disease. New therapies are thus searched as well as biomarkers that would give a better prediction of the risk of relapse. Our study explores the still new field of antisense transcription to define cancer gene lists, and will lead to further works to define predictive markers and/or tailor targeted treatment by antisense oligonucleotides (ASO) ^14^.

This is the first time that a whole transcriptome strand-specific RNA-Seq study focusing on the antisense transcription is performed in paired tumor and healthy mammary tissues. This experimental design allows to detect deregulation of NAT expression that occurs in cancer tissues, and to statistically connect them with the changes of the corresponding coding transcript expression. In particular, we revealed that many positive correlations between NATs and their PCT counterpart were appearing or fading in the tumor, suggesting newly acquired or lost regulations of the protein-coding transcripts in the cancerous tissues. Further functional molecular studies will however be needed to confirm the existence of such regulations of the PCTs by their NATs in the list of cancer gene pairs that were highlighted in this work.

The difference in fold-change between NATs located in genomic alterations and their coding counterparts, with NATs showing higher fold-changes, as well as the significant increase in NAT expression in tumors tend to indicate that NATs may be subject to a particular activating mechanism specific to tumors (Figure 3 A, Table 2).

Moreover, the association of these NATs with survival, which was evaluated through the use of their protein-coding counterparts as proxy in a large independent cohort, shows that the dysregulation observed within the landscape of NATs is not merely a random byproduct of the tumoral process. Analyses were conducted to explore the relationship of the PC and AS genes with known prognostic factors, but no significant results were found. In the same way, enrichment analyses in pathways genes were conducted without any noticeable results.

Several studies have already explored the role of antisense transcription in breast cancer ^15–17^. Grinchuk *et al.* analyzed NAT/PCT pairs that are deregulated in breast cancer in order to define pathways in which they are particularly involved, and they defined NAT/PCT-based prognosis signatures. However Affymetrix microarray datasets were the support of this work, and this technique is not intrinsically strand-specific ^15^. Moreover, mammary normal and tumoral analyzed tissues were not matched. Balbin *et al.* performed a large scale, genome wide, stranded RNA-Seq study on 376 cancers samples with, among them, 60 primary breast cancers ^16^. But as in Grinchuck’s study, tumorous tissues were not matched with healthy ones and in consequence, these studies did not explore, patient by patient, if the NAT/PC expression correlations were already present in the normal tissue, or if they were newly acquired in the tumor. This particularity in our experimental design did allow highlighting the fact that NAT expression is increased in tumorous tissues when compared to their coding counterpart. Indeed, the proportion of NAT reads counts in NAT/PCT pairs is globally raised in tumors when compared to healthy tissues.

As Balbin *et al*. have stated before, at any locus where PCT and NAT are simultaneously transcribed, the PCT is expressed around 1000 times more than the NAT, but we have additionally observed that this difference of expression is lower in tumors than in healthy tissues. We also measured that globally, 10% of the transcripts were coming from the antisense strand in healthy tissues and that this proportion is increased to 13% in tumors (8% were described by Balbin *et al*.). However, some patients present a much higher increase of NAT/PCT proportion in the tumor than others. This heterogeneity in NAT expression deregulation across patients could be used to stratify patients into subgroups of different prognosis. One limitation of our study is the small size and the short follow-up time of our cohort, which did not allow performing such type of analysis.

Our results also confirm the observation by Grinchuk *et al* that the expression correlations between NAT and PCT were different in tumors when compared to unrelated healthy tissues. We refined this observation by using paired tissues of the same patient, and showed that globally these correlations become more significant and more positive in the tumors. Moreover, we highlighted the gene pairs where potential new PCT/NAT expression regulation occurs in cancerous tissues. After having performed a survival analysis with gene expression data from an external cohort (TCGA), it appears that these NAT/PCT gene pairs were also enriched in survival-associated genes, suggesting that the opposite strand transcription regulation might play a role in the breast cancer disease.

Therefore, our report opens a new field of investigation in cancer and indicates that NAT expression is often increased in cancer samples as compared to matched normal tissues. The relevance of this observation for coding gene expression, cancer biology, prognosis and treatment will need to be determined in specific and large cohorts of paired samples.

### Material and Methods

#### Ethical Statement

Tissues were obtained from the Liege University Biobank (N=12) and from the St Vincent Clinic of Rocourt (N=11), Belgium. This study was approved by the local institutional ethical board (2010/229). All aspects of the study comply with the Declaration of Helsinki. Patients of the Liege University Hospital were recruited on the basis of an opt-out methodology. Patients of the St Vincent Clinic of Rocourt were informed of the research work and provided written informed consent.

#### Patients and samples

This retrospective study was performed on 23 cryopreserved cancerous and adjacent healthy tissues from 23 women suffering from estrogen receptor expressing breast cancer. Samples were collected from 2010 to 2014. One patient was excluded because of poor strand-specificity of the RNA-Seq. The clinical and pathological parameters of the patients included in the final analysis were recorded and summarized in Table 1.

A summary of the experimental design is depicted in Figure 1.

#### DNA/RNA/miRNA extraction

DNA, RNA and miRNA were simultaneously extracted using All Prep DNA/RNA/miRNA Universal kit (Qiagen, Belgium) according to the manufacturer protocol. The RNA quality was assessed using a BioAnalyzer (Agilent, Belgium).

#### TruSeq^®^ Stranded Total RNA by Illumina^®^ and next generation sequencing

RNA-Sequencing libraries for 22 breast tumors and paired adjacent tissues were constructed from 500 ng of total RNA, using the TruSeq^®^ Stranded Total RNA kit and Ribo-Zero rRNA Removal kit. A step of chemical fragmentation generated RNA fragment of 180 pb. This step was adapted according to the RNA quality as described in the manufacturer’s protocol. The syntheses of first and second strand of cDNA were performed with random hexamer primers. A 12 cycles PCR was performed to amplify the libraries. The quality and the size of cDNA libraries were assessed using Bioanalyzer Agilent Chip DNA 1000. Only libraries from 290bp to 300pb were used, and 14 pmol final cDNA libraries were loaded on a Illumina HiSeq 2000 apparatus for cluster generation and paired-end sequencing of 2x100 bp, with a mean of 8.26E+09 bases sequenced for each sample (4 samples/flow cell). Kits and apparatus were from Illumina, The Netherlands.

#### CGH array

Array comparative genomic hybridization was performed in the healthy and tumorous tissues of the 22 patients using the Agilent’s 60 K microarray platform (G4827A-031746; Agilent Technologies, Santa Clara, CA, USA) according to the manufacturer’s instructions. The arrays were scanned with SureScan High Resolution Microarray Scanner (Agilent Technologies, Santa Clara, CA, USA). Data and images were imported using the Feature Extraction V.9.5.3.1 Software and results were analyzed with CytoGenomics software v2.5 (Agilent Technologies, Santa Clara, CA, USA). The Aberration Detection Methods 2 algorithm (ADM-2) was used with a cut-off 6.0, followed by a filter to select regions with three or more adjacent probes and a minimum average log2 ratio ± 0.25, was used to detect copy number changes. The quality of each experiment was assessed by the measurement of the derivative log ratio spread with CytoGenomics software v2.0. Genomic positions were based on the UCSC February 2009 human reference sequence (hg19) (NCBI build 37 reference sequence assembly). Filtering of copy number changes was carried out using the BENCHlab CNV software (Cartagenia, Leuven, Belgium).

#### Gene expression quantification by RNA-Sequencing

A quality control of the sequenced reads has been performed with the FastQC software (v. 0.11.2; https://www.bioinformatics.babraham.ac.uk/projects/fastqc/). Sequencing reads were mapped to the Human Genome hg19 GRCh37-75 (Ensembl) using the Star 2.4.1c software ^20^. Mapping quality was assessed with the Picard RnaSeqMetrics tool of the Picard software suite (v. 1.127; http://broadinstitute.github.io/picard/) using default parameters. The results are available in S1-Supplementary Figures. Read counts assignment was performed with the htseq-count tool of the HTSeq software suite (v. 0.6.1) ^21^. Data quality assessment was performed by computing the strand specificity (ratio of sequencing reads mapping to the incorrect strands) of all samples with htseq-count, leading to the removal of one patient with aberrant strand specificity. The DESeq2 software (v. 1.10.1) was used to normalize read counts, estimate dispersion, perform variance stabilizing transformation, and perform independent filtering by using the mean of normalized counts as filter statistic, thereby adjusting the filtering threshold at 33%, following the standard workflow ^22^. Variance-stabilization performance was assessed by producing MA-plots of log2 fold-change versus mean expression with DESeq2. A search for outliers was performed by computing Cook’s distances for every gene and every sample with DESeq2 (S2/S3/S4/S5/S6-Supplementary Figures). A principal component plot was performed to assess the appropriate separation between the 2 sample classes (S7-Supplementary Figures). All aforementioned quality and performance measures yielded acceptable results for all remaining samples.

#### Quantitative RT-qPCR

Reverse transcription was performed using the Reverse Aid H Minus kit (LifeTechnologies, Belgium) from 100 ng of total RNA using random hexamer primers in case of coding genes and using a target specific primers coupled to an unrelated synthetic DNA oligonucleotide in case of NAT.

Quantitative PCR were performed using specific 6-FAM/ZEN/IBFQ probes (IDT, Belgium) with Kapa Probe Fast qPCR Master Mix (Sopachem, Belgium) on a LightCycler 480 apparatus (Roche). In case of coding gene amplification, the primers were designed according to standard procedure. In case of NAT gene amplification, a primer specific to the target NAT and a primer specific to the synthetic oligonucleotide added during the reverse transcription are used to allow a stand-specific amplification.

The relative expression was calculated using the standard curves methods, using β2-microglobuline as endogenous standard.

Primers and probes sequences can be found in the Supplementary Materials and Methods.

#### External dataset used for RNASeq gene expression comparison

Gene expression variations were retrieved from de GEO Dataset GSE65216 (https://www.ncbi.nlm.nih.gov/geo/query/acc.cgi?acc=GSE65216), which is an expression study by micro-array (Affymetrix) of the Maire’s breast cancer cohort ^23^.

#### Definition of the protein-coding/antisense pairs

The list of pairs of protein-coding genes and their corresponding antisense has been generated based on the human genome assembly and gene annotation GRCh37 (release 75) from Ensembl ^24^. To be included in the list, pairs of genes have to fulfill the 3 following conditions: overlapping coordinates; opposite strands; one of the two genes has to have the protein_coding biotype, while the other can have any of the following biotypes: *3prime_overlapping_ncrna, antisense, IG_C_pseudogene, IG_V_pseudogene, lincRNA, misc_RNA, polymorphic_pseudogene, processed_transcript, pseudogene, sense_intronic, sense_overlapping, snoRNA, snRNA*. The reasoning behind including all non-protein_coding biotypes as putative antisenses is that the Ensembl annotation of antisenses is limited to already validated antisenses, and it might thus miss out previously unknown antisenses.

#### NAT/PCT Gene list selection methods

1) DiffCor list: Differential correlation analysis between pairs of protein-coding and antisense transcripts was performed with the DGCA software (v. 1.0.1) ^19^. Pairs of protein coding/antisense genes whose correlation is significantly different between normal and tumor samples (adjusted p-value < 0.05) and whose correlation class differs between tumor and normal samples (ie. we removed the 0/0, +/+, -/- classes) have been selected.

2) NATDiffExp list: Differential expression analysis between all tumor and healthy samples was performed with the DESeq2 software (v. 1.10.1) for all genes, following the standard multi-factor workflow for paired samples. Pairs of protein coding/antisense genes where the antisense was significantly differentially expressed (adjusted p-value < 0.05) between normal and tumor samples have been selected.

3) varRatio list: The read counts ratio variation analysis had been performed as follows: let us define the variation of read counts ratio (varRatio) for each pair of NAT/PCT genes as

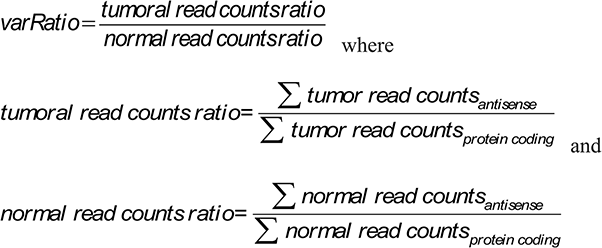

Pairs of NAT/PCT genes corresponding to extreme values of the varRatio distribution have been selected by applying a threshold (mean ± standard deviation) to the log-transformed distribution of the varRatios (S8-Supplementary Figures).

For all three gene list selection methods, pairs of genes where either the protein-coding or the antisense was expressed in less than 7 tumor samples or 7 healthy samples have been discarded.

#### Survival analysis

All protein-coding genes of the 3 gene lists have been tested for association with survival on an external dataset of 1066 RNA-Seq samples from the tumors of female breast cancer patients (Package R TCGA Biolinks; ^25^). Association with survival was recorded when the p-value of a log-rank test was inferior to 0.05. The ratio of genes associated with survival in each list has been compared with the same ratio computed with a list of randomly selected protein-coding genes.

## Acknowledgments

We thank the GIGA-genotranscriptomic-platform with special attention to Wouter Coppieters, the team of medical oncologists and data managers of the Medical Oncology Department, and the Biothèque of CHU Liège. We also thank Jérome Thiry, Bouchra Boujemla and Christophe Poulet.

